# Efficient reprogramming of the heavy-chain CDR3 regions of a human antibody repertoire

**DOI:** 10.1101/2021.04.01.437943

**Authors:** Tianling Ou, Wenhui He, Brian D. Quinlan, Yan Guo, Pabalu Karunadharma, Hajeung Park, Meredith E. Davis-Gardner, Mai H. Tran, Yiming Yin, Xia Zhang, Haimin Wang, Guocai Zhong, Michael Farzan

## Abstract

B cells have been engineered *ex vivo* to express an HIV-1 broadly neutralizing antibody (bNAb). B-cell reprograming may be scientifically and therapeutically useful, but current approaches limit B-cell repertoire diversity and disrupt the organization of the heavy-chain locus. A more diverse and physiologic B-cell repertoire targeting a key HIV-1 epitope could facilitate evaluation of vaccines designed to elicit bNAbs, help identity more potent and bioavailable bNAb variants, or directly enhance viral control *in vivo*. Here we address the challenges of generating such a repertoire by replacing the heavy-chain CDR3 (HCDR3) regions of primary human B cells. To do so, we identified and utilized an uncharacterized Cas12a ortholog that recognizes PAM motifs present in human JH genes. We also optimized the design of 200 nucleotide homology-directed repair templates (HDRT) by minimizing the required 3’-5’ deletion of the HDRT-complementary strand. Using these techniques, we edited primary human B cells to express a hemagglutinin epitope tag and the HCDR3 regions of the bNAbs PG9 and CH01. Those edited with bNAb HCDR3 efficiently bound trimeric HIV-1 antigens, implying they could affinity mature in vivo in response to the same antigens. This approach generates diverse B-cell repertoires recognizing a key HIV-1 neutralizing epitope.

## INTRODUCTION

Traditional vaccination approaches do not elicit broadly neutralizing antibodies (bNAbs) that target conserved epitopes of HIV-1 envelope glycoprotein trimer (Env).^1-4^ Human precursor B-cell receptors (BCRs) that can develop into bNAbs are rare,^5, 6^ and mature bNAbs have properties that are difficult to access through antibody maturation.^1, 7, 8^ A number of groups have begun to explore an alternative to conventional vaccines in which B cells themselves are reprogrammed.^9-13^ This approach employs CRISPR-mediated editing of the BCR loci so that the edited B-cell expresses a mature HIV-1 bNAb. In addition to its long-term potential for reprograming human immune responses, BCR editing can be applied more immediately to generate animal models useful for assessing vaccination strategies, and for developing more potent and bioavailable bNAb variants.

The BCR includes a membrane-bound heavy chain (H) covalently associated with a light chain (L). Both chains are composed of a variable and a constant region. The heavy-chain variable domain is formed by a process of VDJ recombination of the immunoglobulin heavy chain (IgH) gene. In humans, one of the 38-46 functional variable (VH) genes recombines with one of 23 diversity (DH) and one of 6 joining (JH) genes.^14^ The recombination process also introduces diversity at the junctions of VH, DH, and JH genes through removal and addition of nucleotides. The light-chain variable domain is formed similarly by VJ recombination of the IgL gene. The naïve B-cell repertoire thus reflects extensive combinatorial diversity.^15-18^ This diversity is further amplified after antigen exposure. B cells undergo somatic hypermutation (SHM) as they compete for access to antigen in the lymph-node germinal centers, a process resulting in affinity maturation of the BCR.^19^

The combinatorial diversity of the B-cell repertoire complicates efforts to reprogram BCR. To date, investigators have bypassed this challenge by targeting an unvarying intron between the recombined variable region and the IgM constant region (Cµ).^9, 11-13^ This strategy introduces a single cassette encoding an exogenous promoter and bNAb heavy- and light-chain sequences into this heavy-chain intron. By design, these constructs halt expression of the native variable heavy chain. Expression of the native B-cell light chain is usually also prevented through various mechanisms. While powerful and convenient, this approach eliminates combinatorial diversity and relies solely on SHM to broaden the HIV-1 neutralizing response. In addition, it introduces several less physiologic elements including novel locations for both variable genes, use of exogenous promoter, and some architectural differences between the expressed bNAb-like construct and native antibodies. These limitations may be especially important if edited B cells need to adapt efficiently to a diverse HIV-1 reservoir,^20, 21^ or when the edited repertoire is used to study B cell biology.^22^

Here we develop a complementary approach in which sequence encoding a bNAb HCDR3 is introduced into a diverse BCR repertoire at its native location. This approach is useful with antibodies that are highly dependent on the HCDR3 to bind antigen, including members of an exceptionally potent class of bNAbs that recognize the Env apex/V2-glycan epitope.^23, 24^ However, retaining combinatorial diversity in a natural B-cell setting poses several challenges. First, long homology arms of a homology-dependent repair (HDR) template (HDRT) can overwrite the V-encoded region of a heavy chain. Second, the region 5’ of the HCDR3-encoding region is necessarily diverse, and thus editing can be variably efficient due to mismatch of a homology arm with its chromosomal complement. Third, introducing exogenous HCDR3 requires deletion of chromosomal material of unknown length and content, rather than simply insertion of sequence, or direct replacement of a known sequence. To address these challenges, we identified a previously uncharacterized Cas12a variant^25^ that efficiently recognizes a specific four-nucleotide protospacer adjacent motif (PAM) present in the 3’ region of the most commonly used JH genes in humans. We also optimized the use of 200-nucleotide (nt) single-stranded HDRT with short (50 nt) homology arms, demonstrating in the process that editing efficiency is primarily determined by the length of a 3’ mismatch tail rather than the relationship of the HDRT to transcription direction (sense or antisense) or to the target strand of the CRISPR guide RNA (gRNA). With these procedures, we altered the specificity of primary human B cells by editing their HCDR3 regions to bind HIV-1 Env, while retaining the original diversity of the VH repertoire. These studies demonstrate the feasibility of an alternative approach to human B-cell reprogramming.

## RESULTS

### Targeting a conserved region of the immunoglobulin heavy chain locus with a Cas12a ortholog

A major challenge of precisely replacing the HCDR3-encoding region of a diverse primary B cell population is the variability of the mature IgH locus. This variability arises from the random combinations of V, D, and J segments which are joined imprecisely and unpredictably.^16^ It complicates two processes necessary for CRISPR-mediated editing of the B-cell locus, namely the selection of a guide RNA (gRNA) that must complement a 20-24 nucleotide genome sequence, and the design of HDRT whose 5’ and 3’ homology arms must complement even longer genomic regions.^26-28^

We began by designing a gRNA that recognizes a large proportion of BCR and targets genome cleavage to site of insertion, where it is most efficient. The HCDR3 is encoded by of the 3’ end of a VH gene, a DH gene, and the 5’ end of a JH gene (**Figure 1A**). There are six human JH segments, and a JH4 alone participates in 50% of productive human VDJ-recombination events.^29, 30^ Due to junctional diversity, the 3’ JH region is conserved, but the 5’ is less predictable. However, the 3’ of JH4 did not contain any canonical Cas12a PAM^31^ sequences (TTTV), and the available Cas9 PAM (NGG)^32^ mediates cleavage too distal from the site of insertion. Instead, two potential non-canonical Cas12a PAM sites, GTTC and TTCC,^33^ were well positioned to facilitate gRNA recognition of conserved JH regions while cleaving where a new HCDR3 would be inserted. We therefore characterized a number of Cas12a orthologs for their ability to recognize these divergent PAM sites. To do so, we tested several uncharacterized Cas12a orthologs^25^ in the human B-cell line Jeko-1. Jeko-1 cells were transfected with plasmids encoding BsCas12a, TsCas12a, Mb2Cas12a, or Mb3Cas12a along with plasmids encoding gRNAs adjacent to the GTTC and TTCC PAM regions. DNA cleavage within the HCDR3 frequently results in error-prone nonhomologous end joining (NHEJ) that eliminates expression of the Jeko-1 BCR.^34^ Thus loss of IgM expression indicates a successful double-strand break. Among the Cas12a orthologs tested, Mb2Cas12a most efficiently cleaved the Jeko-1 JH4 region initiated with GTTC and TTCC, with the highest efficiency (18.9%) observed when the GTTC PAM was targeted (**Figure 1B**).

**Figure 1.**
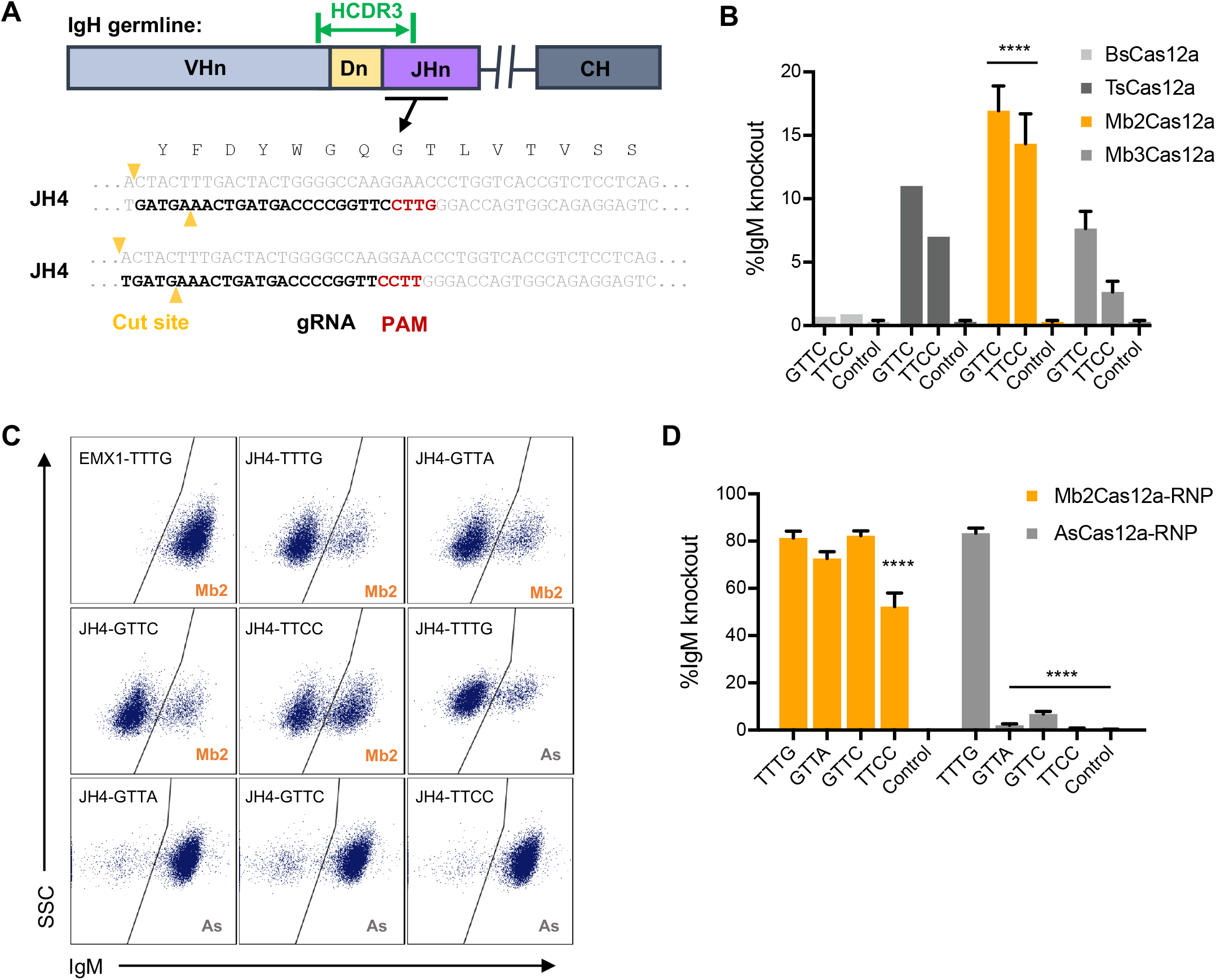
Targeting the conserved region of JH4 gene requires a Cas12a ortholog recognizing non-canonical PAMs. (**A**) A representation of the coding region of an antibody heavy-chain variable region is presented. As indicated, the HCDR3 (green) is encoded by the 3’ of a recombined V gene, a D gene, and the 5’ of a J-chain. To insert a common HCDR3 into a diverse population of BCR, the guide RNA (gRNA) of a CRISPR effector protein must complement a conserved HC region at the 3’ of the recombined J-gene, while cleaving a more variable region near the site of HCDR3 insertion. Note that, unlike Cas9, Cas12a cleaves distally from its PAM and seed regions. The preferred PAM recognition sequence of commonly studied Cas12a orthologs is TTTV. However, as shown, JH4, the most frequently used JH gene in all species, contains optimally located GTTC and TTCC PAM sequences, located 3’ of the HCDR3-encoding sequence but oriented Cas12a cleavage within the this sequence. This PAM, sequence of the gRNA, and the Cas12a cut sites are indicated. (**B**) To identify a Cas12a ortholog efficient at cleaving these non-canonical PAM motifs, the human B-cell line Jeko-1 was transfected with plasmids encoding BsCas12a, TsCas12a, Mb2Cas12a, or Mb3Cas12a. Targeting efficiency was measured by flow cytometry as loss of IgM expression. Among these Cas12a orthologs, Mb2Cas12 most efficiently cleaved the J-chain region initiated with GTTC and TTCC (orange). Error bars indicted range of two independent experiments, and asterisks indicate statistical significance relative to controls. Statistical difference were determined by non-paired Students t-test, (****, p<0.0001). (**C**) Mb2Cas12 RNP were compared with commercial AsCas12a RNP for their ability cleave four distinct regions in the HCDR3-encoding region of Jeko-1 cells. Loss of IgM expression indicates successful introduction of a double-strand break and inexact NHEJ. (**D**) Results of three experiments similar to that shown in panel C. Error bars indicate standard error (SEM). Asterisks indicted significant differences from the canonical TTTG PAM (Mb2Cas12a-RNP or AsCas12a, respectively). Statistical difference were determined by non-paired Students t-test, (****, p<0.0001).

Ribonucleoprotein (RNP) forms of CRISPR effector proteins, electroporated into cells, are typically more efficient than plasmids expressing the same protein.^28, 35^ We accordingly produced Mb2Cas12a RNP and compared its editing efficiency with a commercial AsCas12a RNP in Jeko-1 cells, again as determined through loss of IgM expression. These RNP cleaved a canonical Cas12a TTTG with comparable efficiency but Mb2Cas12a cleaved three non-canonical PAM regions more efficiently (**Figures 1C and 1D**), demonstrating that Mb2Cas12a has a broad PAM specificity and efficiently edits the HCDR3 region of Jeko-1 cells. Notably, Mb2Cas12a RNP efficiently cleaved JH4 when initiated with a GTTC PAM, and this PAM region is conserved in the JH-genes of both humans and rodents.

### Optimization of gene editing using single-stranded HDR templates

We also optimized the design of HDRT used to replace a native HCDR3 region. Again, the underlying diversity of the recombined heavy-chain limited our options. Most importantly, the HDRT homology arms needed to remain short to maximize complementarity to the 3’ VH region. We therefore optimized a strategy based on short single-stranded DNA (ssDNA) HDRT with 50-nucleotide homology arms by monitoring the efficiency with which a hemagglutinin (HA) tag could replace the Jeko-1 HCDR3 region. Specifically, we compared sense and anti-sense forms of two distinct HDRT, each with different length linkers bounding the HA tag, and cleavage at four distinct Mb2Cas12a sites and four proximal SpCas9 sites (**Figure 2A and Figure S1**). Note that some of these sites are unique to the Jeko-1 HCDR3 region, and are therefore not generalizable to primary B cells. Knock-in efficiency was determined by flow cytometry with fluorescently labelled anti-HA antibodies (**Figure 2B**). We observed that on average, with four different cut sites and four distinct HDRT, Mb2Cas12a and SpCas9 edited with comparable efficiencies (**Figure 2C**). We analyzed these same data by comparing a number of parameters proposed to impact editing efficiencies in other systems.^27, 28, 36, 37^ However, no significant differences were observed when sense or anti-sense strand HDRT were used (**Figure 2D**), and only a modest difference for Mb2Cas2, but not for SpCas9, was observed when target (complementary to gRNA) or non-target strand HDRT was used (**Figure 2E**). These data suggest that our system is distinct from those of previous studies, perhaps because both DNA deletion and insertion are required to replace an HCDR3.

**Figure 2.**
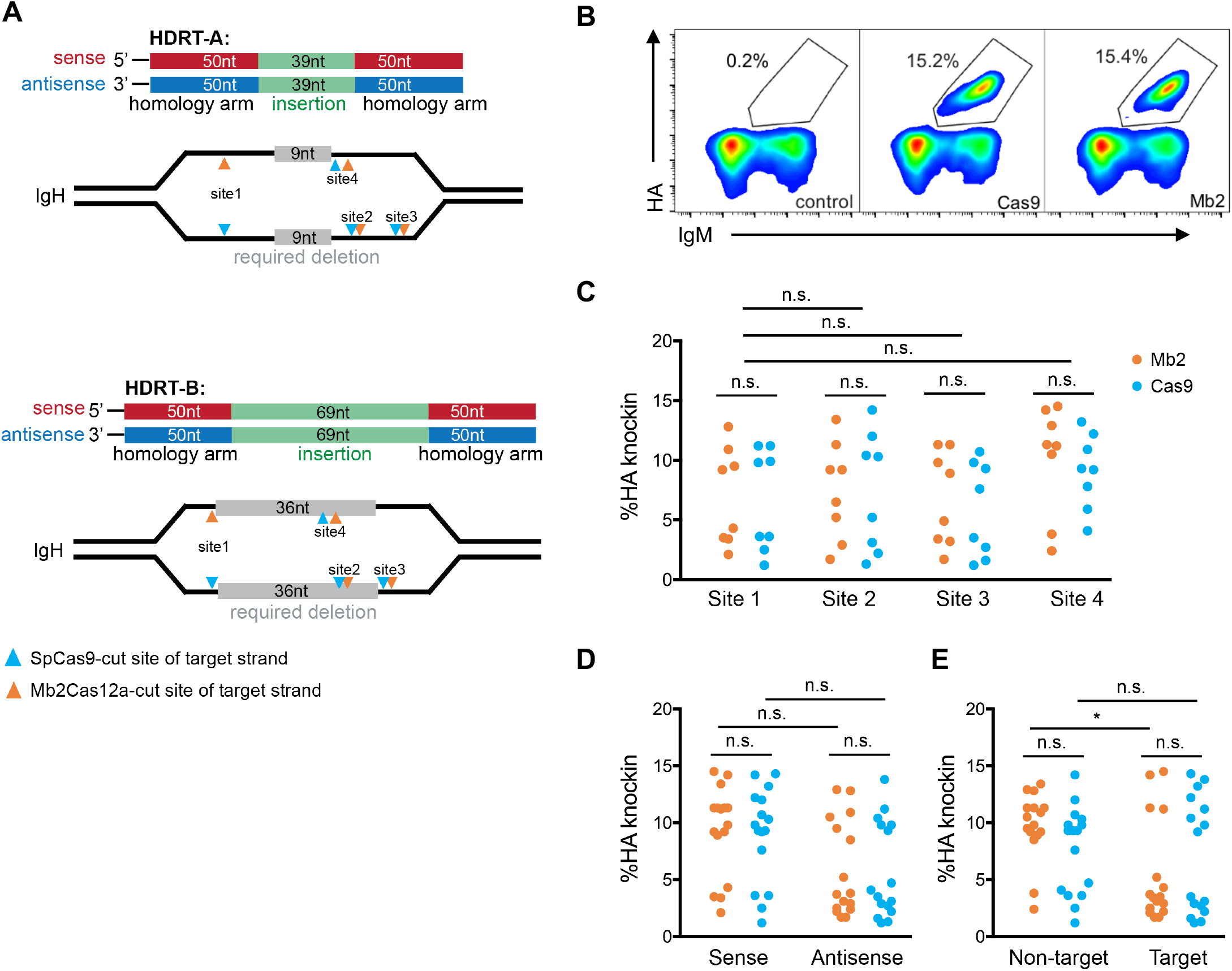
Optimization of ssDNA templates for Mb2Cas12a-mediated editing the HCDR3-encoding region of a human B cell line. (**A**) A diagram representing four HDR templates (HDRT) used in panels B-E. Specifically, sense and anti-sense forms of HDRT-A, were used to replace a 9-nucleotide (nt) region (grey) with 39-nt insert (green), and both forms of HDRT-B were used to replace a 36-nt region with a 69-nt region. 50-nt homology arms of the sense and antisense forms are represented in red and blue, respectively. SpCas9 (cyan) and Mb2Cas12 (orange) cleavage sites of the target strand (complementary to grNA) are indicated by arrows. Note that paired Cas9 and Cas12a cleavage sites are separated by at most five nucleotides. (**B**) A representative example of an experiment used to generate panels C-E in which editing efficiency of MbCas12A or SpCas9 RNP is monitored through recognition of an HA tag introduced into the HCDR3 of the Jeko-1 cell BCR by flow cytometry. Control cells were electroporated with Mb2Cas12a RNP without an HDRT. (**C**) A comparison of Mb2Cas12a (Mb2) and SpCas9 (Cas9) knock-in efficiencies, measured as described in panel B, for all four sites shown in panel A. Differences between Mb2 and Cas9, and among the four sites, are not significant (n.s.). The same data generated for panel C was replotted according to whether the sense or anti-sense HDRT were used (**D**), or whether the HDRT complemented the gRNA target or non-target strand. (**E**) Non-target strand is the PAM containing strand, and the target strand is the strand annealed to gRNA. Again, as indicated, most differences were not significant. However, the HDRT complementary to the Mb2Cas12a gRNA target strand were slightly more efficient than those complementary to the non-target strand (p=0.027). Dots in (**C**)-(**E**) represent pooled data from two independent experiments. Statistical significance was calculated by one-way ANOVA with Tukey’s multiple comparison test.

**Figure 3A** presents a model of a system in which deletion of chromosomal DNA and insertion of novel sequence are both necessary for successful editing. After CRISPR-mediated formation of a double-strand break, HDR is initiated by 5’ to 3’ resection of DNA, exposing two 3’ ends, one of which can anneal to a single-stranded HDRT homology arm. However, at least one 3’ end necessarily includes sequence that must be deleted 3’ to 5’, either on the HDRT-complementary strand, creating a 3’ mismatch tail, on the opposing strand, or on both. As shown in **Figures 3B and 3C**, the length of this 3’mismatch tail strongly predicts editing efficiency in this system regardless of whether the double-stranded break was mediated by Mb2Cas12a or SpCas9. Specifically, editing is significantly more efficient when mismatch tails are shorter than 10 nucleotides, presumably because longer tails prevent polymerase priming and templated extension of the 3’ arm. We presume that this principle can be extended to other systems in which a templated sequence must replace a chromosomal region by HDR.

**Figure 3.**
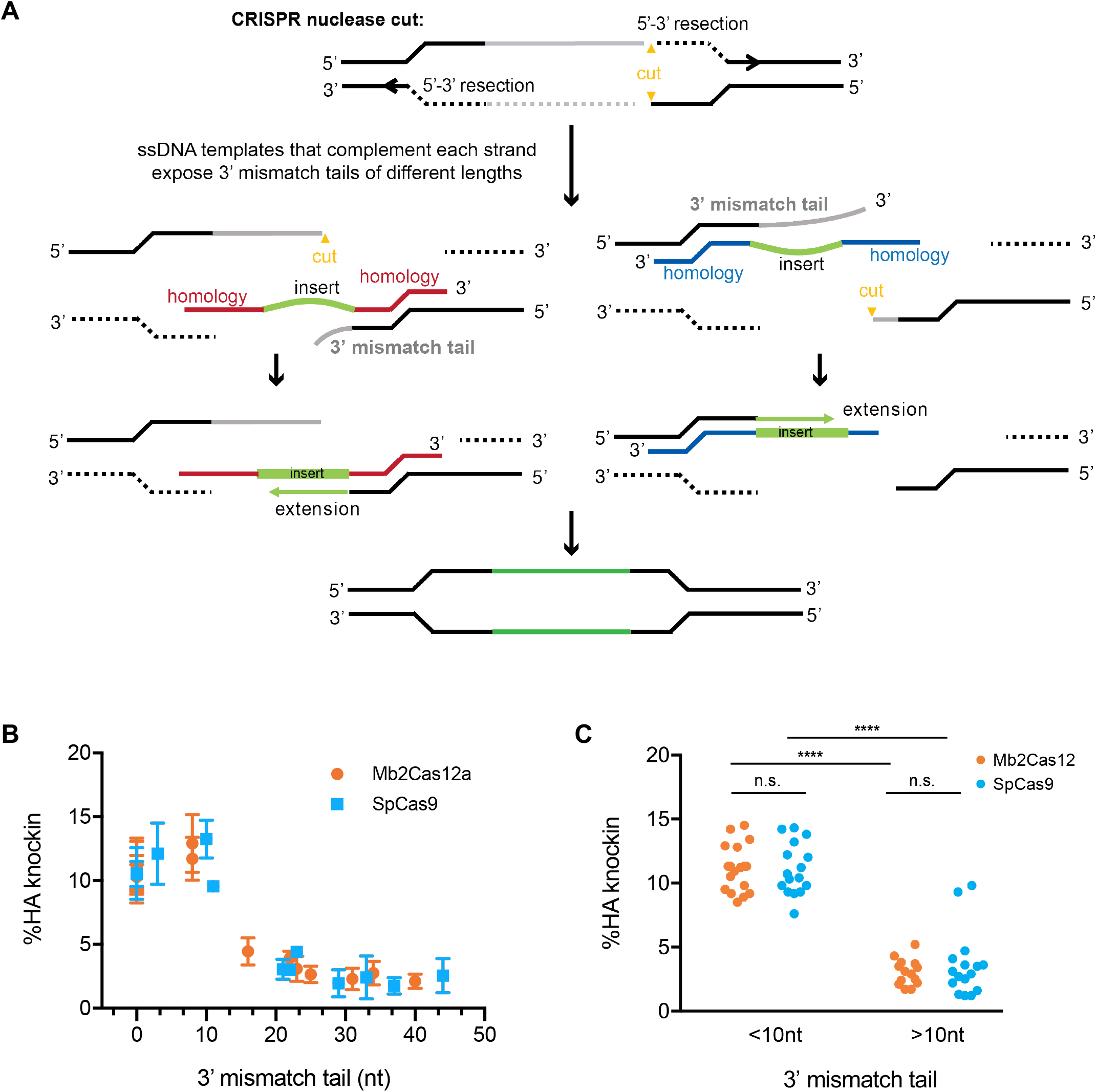
The length of the 3’ mismatch tail determines replacement efficiency with short single-stranded HDRT. (**A**) A model showing where a 3’ mismatch tail occurs. A cut site (yellow) is introduced into a region of the gene targeted for replacement (grey), asymmetrically dividing this region. Efficient 5’ to 3’ resection exposes two 3’ ends. An HDRT can complement a strand with a short (left figures) or long 3’-mismatch tail (right figures), which must be removed before the remaining 3’ end can be extended to complement the HDRT insert region and its distal homology arm. We propose that the removal of this 3’ mismatch tail is a rate-limiting step determining editing efficiency when genomic sequences are replaced. (**B**) The predicted length of the 3’-mismatch tail in experiments presented in Figure 2 are plotted against the efficiency with which an HA-tag is introduced into the HCDR3 region, as determined by flow cytometry. Error bar indication SD from two independent experiments. (**C**) A comparison of editing efficiency between those with short (<10 nt) or long (>10 nt) 3’ mismatch tails. Editing by SpCas9 or Mb2Cas12a is significantly more efficient with short 3’ mismatch tails, as determined by one-way ANOVA with Tukey’s multiple comparison test (p<0.0001). Dots represent pool data from two independent experiments.

### Reprogramming B cell specificity towards HIV through HCDR3 replacement

Using the strategies described above, we then designed HDRT that could replace the Jeko-1 HCDR3 with those of PG9 or PG16 (**Figure 4A**), two potent HIV bNAbs directed against a V2 apex of the HIV-1 Env trimer.^38, 39^ To monitor the successful introduction of these HCDR3 in contexts where the resulting BCR does not bind soluble native-like HIV-1 Env trimer (SOSIP)^40, 41^, we employed an antibody, PSG2,^42^ that recognizes sulfated tyrosines present at the tips of the PG9 and PG16 HCDR3. In addition, we monitored HIV-1 Env binding with two reagents, a SOSIP protein derived from the HIV-1 isolate BG505, and the multivalent nanoparticle (E2p)^43^ based on the same BG505 HIV-1 isolate. Jeko-1 cells were edited with HDRT-PG9-CVR and HDRT-PG9-CAR to express two forms of the PG9 HCDR3, distinguished by an alanine or valine immediately adjacent to the HCDR3-initiating cysteine. Cells edited with either HDRT could be recognized by all three binding reagents, indicating that the resulting BCR could bind the BG505 Env (**Figures 4B and C**). In contrast, Jeko-1 cells edited to express the PG16 HCDR3 were recognized only by PSG2, indicating that editing was efficient, but that the resulting BCR did not bind HIV-1 Env. As expected, none of these three antigens bound Jeko-1 cells in which a control HDRT, introducing an HA-tag into the HCDR3, was employed.

**Figure 4.**
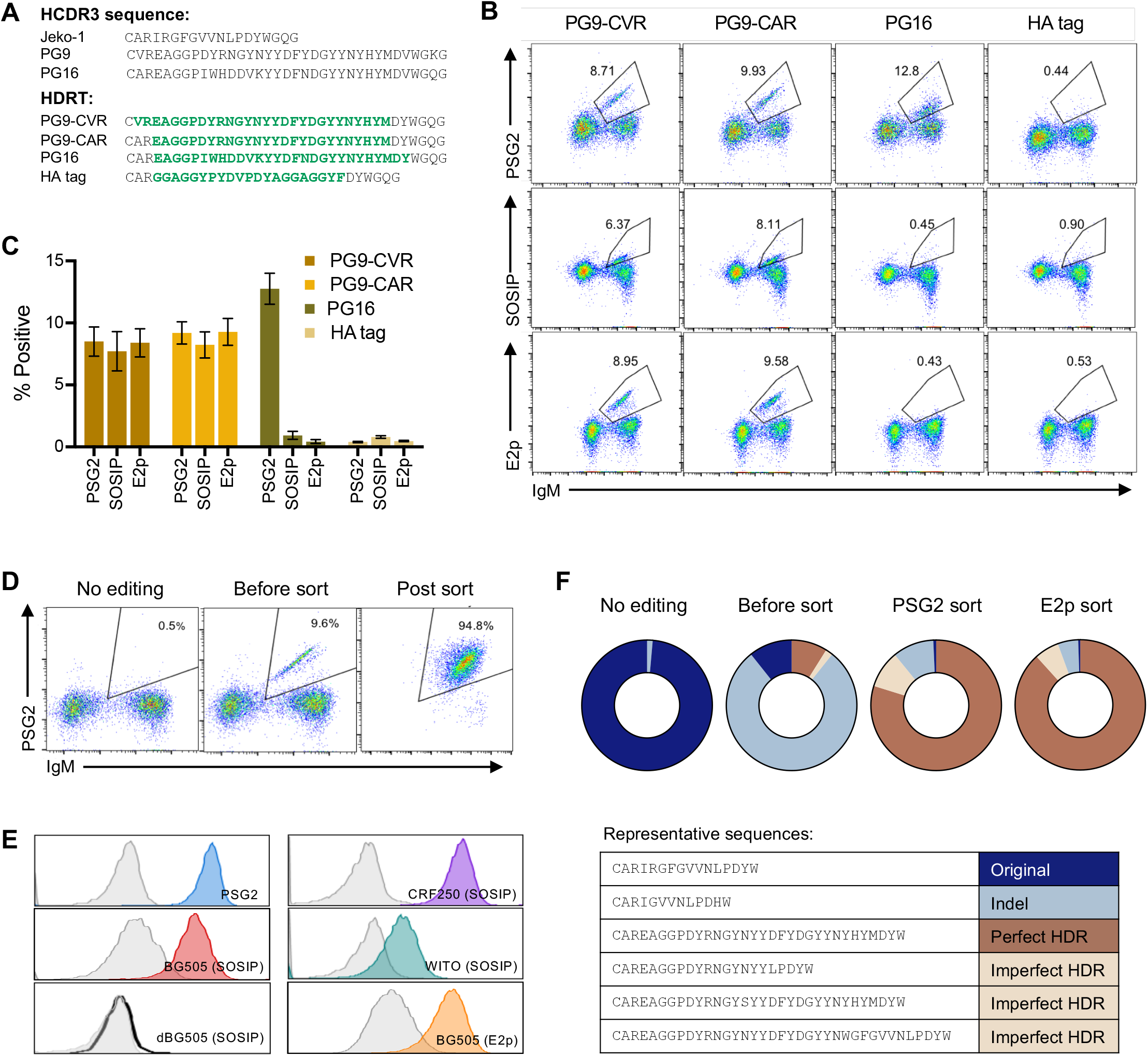
The BCR specificity of Jeko-1 cells can be reprogrammed with a novel HCDR3. (**A**) The amino-acid sequence of the native Jeko-1 cell HCDR3 region and those of the HIV-1 neutralizing antibodies PG9 and PG16 are shown. In addition the amino-acid translations of four HDRT used in the subsequent panels are represented in green, in the context of the remaining Jeko-1 region. (**B**) Mb2Cas12a RNP targeting the GTTC PAM of Site 4 in Jeko-1 cells shown in Figure 2B were co-electroporated with the indicated HDRT. Editing efficiency was monitored on the vertical axis by flow cytometry with fluorescently labeled PSG2, an antibody that recognizes sulfotyrosines within the PG9 and PG16 HCDR3 region, a similarly labeled HIV SOSIP or E2p. The horizontal axis indicates IgM expression, and its loss indicates imprecise NHEJ after Mb2Cas12a-mediated cleavage. Note that introduction of a PG16 HCDR3 was efficient, as indicated by PSG2 recognition, but unlike the PG9 HCDR3, it did not bind the Env trimer. Cells edited to express an HA tag did not bind any reagent. SOSIP proteins were derived from the BG505 HIV-1 isolate. (**C**) A summary of three independent experiments similar to that shown in panel B. flow cytometric studies used to generate panel B. Error bars indicate SD. (**D**) Jeko-1 edited with PG9-CAR HDRT were enriched by FACS with the anti-sulfotyrosine antibody PSG2. (**E**) Cells enriched in panel D were analyzed two weeks later by flow cytometry for their ability to bind PSG2, a BG505-based nanoparticle (BG505-E2p), SOSIP trimers derived from the indicated HIV-1 isolate, or an V2 apex negative mutant (dBG505-SOSIP). Grey control indicates wild-type Jeko-1 cells. (**F**) Unedited Jeko-1 cells and those edited with PG9-CAR HDRT without sorting, or sorted with PSG2 or with E2p, were analyzed by next-generation sequencing (NGS) of the VDJ region. Sequences were divided into four categories, depending on whether the edited sequence matched exactly the HDRT (Perfect HDR), whether HDRT sequence was visible but modified (Imperfect HDR), whether the original Jeko-1 HCDR3 region was intact (Original), or whether this region was modified by NHEJ as indicated by the presence of insertions or deletions (Indel). Representative examples of each category are shown below the charts.

We further characterized Jeko-1 cells edited with HDRT-PG9-CAR by enriching edited cells by FACS with the PSG2 antibody (**Figure 4D**). Sorted cells were then analyzed by flow cytometry for their ability to interact with PSG2, SOSIP variants derived from three HIV-1 isolates and from a negative mutant, and BG505-E2p (**Figure 4E**). Each of these reagents bound PG9-HCDR3-edited cells efficiently, with CRF250 SOSIP proteins binding most efficiently and therefore used in subsequent experiments. In parallel, NGS analysis was performed on the HCDR3 region of unedited Jeko-1 cells, cells edited with HDRT-PG9-CAR before they were sorted, and the same cells sorted with either PSG2 or the E2p nanoparticle presenting multiple BG505 proteins. HCDR3 sequences were divided based on whether HDR was successful and whether the introduced sequence exactly matched that in the HDRT (**Figure 4F**). We observed that before sorting, the original Jeko-1 HCDR3 bearing indels reflecting NHEJ predominated. After sorting, HCDR3 that matched the HDRT predominated. Collectively, the data shown in **Figure 4** indicate that the HCDR3 of Jeko-1 cells can be replaced by that of PG9 to generate a BCR that efficiently binds multiple HIV-1 SOSIP proteins.

### Using consensus sequences of multiple VH families to edit primary human B-cells

The preceding studies showed that Mb2Cas12a could cleave a conserved region of the JH4 gene useful for introducing an exogenous HCDR3 sequence, that editing with sense-strand HDRT in this setting is optimal because it minimizes the length the 3’ mismatch tail, and that the PG9 HCDR3 could function with the divergent Jeko-1 heavy-and light-chain to bind multiple HIV-1 Env trimers. However, primary human B-cells pose an additional challenge: in contrast to Jeko-1 cells, the VH-gene sequences of primary cells are variable and unpredictable. This difficulty complicates the design of the 5’ homology arm, which must complement the 3’ region of the VH gene. Alignment of the 3’ regions of the most commonly used VH gene families, namely VH1, VH3, and VH4 (**Figure S2**) revealed that a good deal of interfamily diversity, but showed that intrafamily diversity was limited among the 3’ nucleotides. We accordingly evaluated HDRT with 5’ homology arms based on consensus sequences for each of these VH families in primary human B cells. These cells were isolated from peripheral blood and activated by an anti-CD180 antibody^11^ for 48 hours before electroporation with Mb2Cas12a RNP along with different HDRTs. Editing efficiency was measured 48 hours post electroporation by flow cytometry using the anti-sulfotyrosine antibody PSG2 (**Figure 5A**). Two different fluorophores were used to label PSG2 to eliminate non-specific binding from either fluorophore. As negative controls, primary human primary B cells were activated in the same way, but they were not electroporated (null), or electroporated with Mb2Cas12a RNP and an HDRT that was not homologous to any sequence in the human genome. The HDRT homologous to VH1 had relatively lower efficiency than to the other two families, largely due to the lower VH1 usage frequency in mature human B cells (**Figure 5B**). A equal mixture of three HDRTs (those of VH1, VH3, and VH4) edited more cells than any individual HDRT, suggesting a diverse pool of B cells could be targeted simultaneously. NGS performed on two sets of B cells from different donors edited with the mixed HDRT, and the frequency of successful in-frame editing reflected the efficiencies of the individual HDRT (**Figure 5C**). These data show that, using consensus HDRT homology arms, approximately 1% of primary human B cells can be edited to express the PG9 HCDR3.

**Figure 5.**
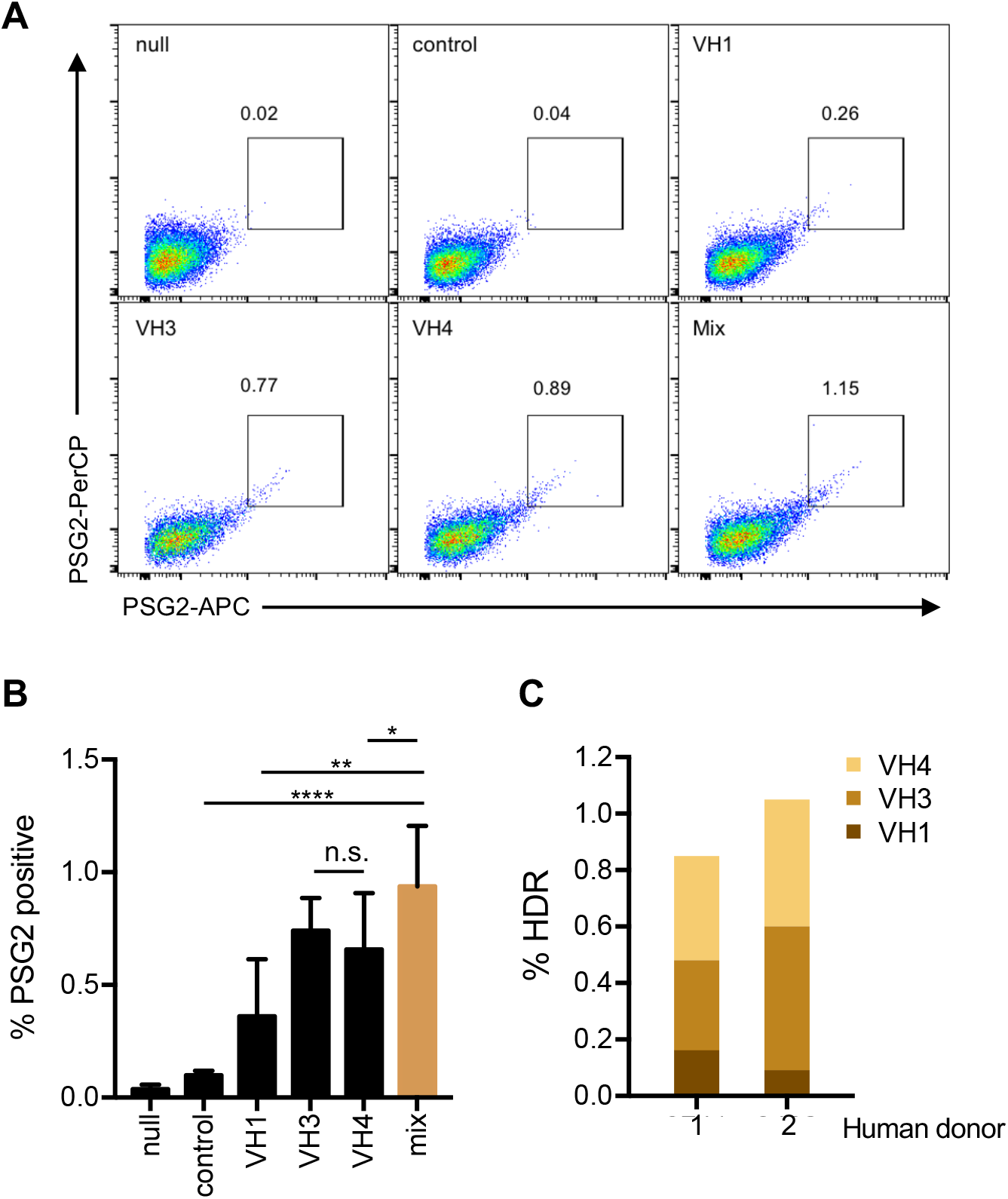
Editing primary human B-cells with HDRT recognizing consensus sequences of multiple VH families. (**A**) A panel of PG9-CAR HDRT with homology arms complementary to JH4 and to consensus VH1-,VH3-, and VH4-family sequences were evaluated for their ability to edited primary human B cells. Cells electroporated with Mb2Cas12a RNP and PG9-CAR HDRT were analyzed by flow cytometry with the anti-sulfotyrosine antibody PSG2 modified with two distinct fluorophores to eliminate non-specific binding from either fluorophore, (**B**) A summary of results from experiments similar to that shown in panel A, using primary B cells from three independent donors. Note that a mixture of three HDRT edited more cells than any individual HDRT. Null indicates that cells were not electroporated and control indicates cells electroporated with Mb2Cas12a RNP and an HDRT that is not homologous to any sequence in the human genome. Mix indicates cells electroporated with RNP and an equimolar mixture of HDRT with VH1-, VH3-and VH4-specific homology arms. Error bars indicted range of three independent experiments, and asterisks indicate statistical significance calculated by one-way ANOVA with Tukey’s multiple comparison test (*, p<0.5; **, p<0.01; ****, p<0.0001). (**C**) NGS analysis of primary B cells from two human donors, quantified as described in Figure 4F except that the VH-family of edited cells was also counted.

### The diversity and specificity of HCDR3-edited primary human B-cell repertoires

To determine if HCDR3-edited BCR acquired their reprogrammed specificity and retained their VH diversity, edited B cells were expanded for a week and then sorted with an appropriate antigen. To evaluate their specificity and diversity, we tested HDRT encoding the PG9 HCDR3, the HCDR3 of the HIV-1 bNAb CH01, or an HA tag (**Figure 6**). As in **Figure 5**, Mb2Cas12a RNP were used to cleave the JH4 region, and mixtures of three HDRT, recognizing consensus 3’ VH1, VH3, or VH4 sequences, directed the insertion of the novel HCDR3. Edited cells were sorted with an anti-HA antibody (**Figure 6A**) or the CRF250 SOSIP trimer derived from the CRF_AG_250 isolate (**Figures 6B and C**). Heavy-chain sequences were analyzed by NGS before (blue) and after (red) sorting. As anticipated, sorting changed the frequency of successfully edited B cells. Critically, in each case, multiple VH1, VH3, and VH4 genes continued to be represented after sorting, suggesting that the combinatorial diversity of the repertoire could be preserved after introducing the PG9 HCDR3 into primary B cells. To confirm that this HCDR3 could function when expressed with multiple variable genes, we expressed in HEK293T cells antibody variants with the PG9 HCDR3 and light-chain, but encoded with several distinct VH genes. We focused on VH1-, VH3-, and VH4-family genes that generated a high signal after sorting, and included VH3-33, the original PG9 variable gene, for comparison. All five VH genes generated antibodies that could detectably express and bind a SOSIP trimer, with BCR expressed from the VH3-30 sequence binding most efficiently (**Figures 6D and E**). Similarly soluble forms of these antibodies could neutralize at least one HIV-1 isolate (**Figure 6F**). Notably, antibodies generated from the VH3-30 gene bound SOSIP trimers and neutralized HIV-1 more efficiently than those based on VH3-33, indicating that VH3-30 and perhaps several other VH genes could serve as alternative starting points for PG9-like antibodies. Collectively, Fig. 6 shows that diverse human B cells can be edited to express the PG9 HCDR3, conferring on many of these cells the ability to bind a SOSIP trimer and express antibody primed to neutralize multiple HIV-1 isolates.

**Figure 6.**
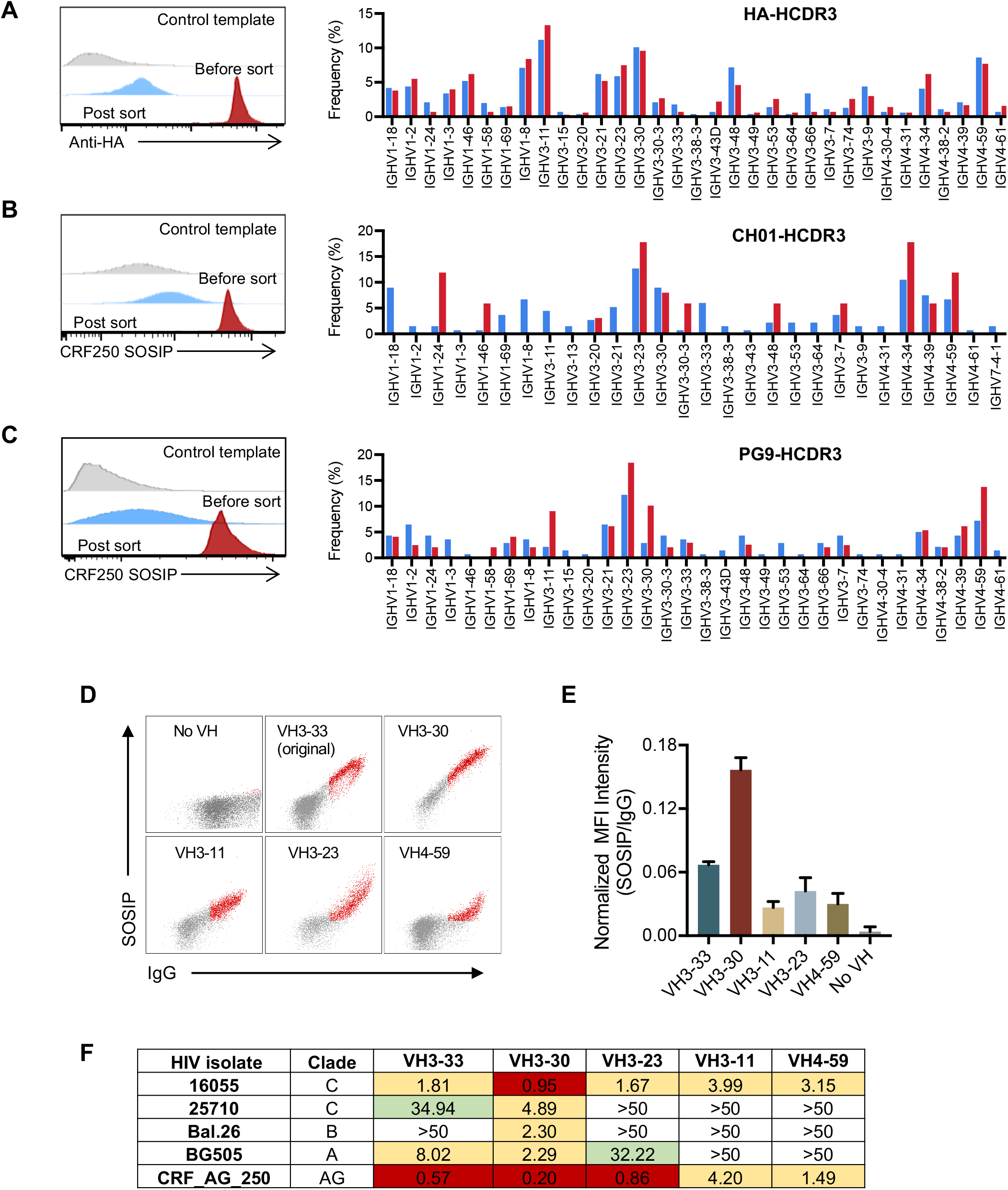
Reprogrammed primary human B cells retain V-gene diversity. Primary cells were electroporated with Mb2Cas12a RNP and HDRT encoding an HA tag (**A**) or the HCDR3 regions of the HIV-1 neutralizing antibodies CH01 (**B**) and PG9 (**C**), with the same mixture of homology arms as those used in Figure 5. Cells were sorted with an anti-HA antibody (HA tag, panel A) or a SOSIP trimer derived from the CRF_AG_250 isolate (panels B and C). Edited cells were analyzed by NGS before and after sorting, and the frequency of each VH1-,VH3-, and VH4-family gene was measured. Flow cytometry histograms displays one of two experiments with similar results, and bar graphs indicate the mean of those two experiments. (**D**) Antibodies composed the heavy-chains expressed from the indicated VH genes enriched in panel C or that of PG9, the PG9 HCDR3, a transmembrane domain, and the native PG9 light chain were expressed on the surface of 293T cells and analyzed by flow cytometry. One of two representative experiments is presented. Mature indicates expression of the original PG9 antibody. (**E**) The mean of two experiments shown in panel D is presented. (**F**) The IC_50_ values of soluble forms of the antibodies characterized in panel D against indicated HIV-1 isolates is represented.

## DISCUSSION

Transgenic mice engineered to expressed human variable-chain sequences of bNAbs and inferred germline forms of these bNAbs have been used extensively to study bNAb maturation in response to HIV-1 antigens.^44-46^ These mice were developed primarily to study vaccination strategies, but they could potentially be used to improve the breadth and potency of bNAbs as well. The advent of CRISPR technologies enables *ex vivo* editing of mature B cells, and adoptive transfer edited B cells into a new murine host.^9, 11-13^ CRISPR-mediated editing of B cells is more rapid and versatile than developing a transgenic mice, but a more limited subset of B cells express the bNAb of interest. These cells can nonetheless be amplified through vaccination, and they undergo class switching and SHM. As this technology advances, reprogramming of mature naïve B cells could replace transgenic mice as a means of evaluating and optimizing vaccine protocols. This approach could also be developed as an *in vivo* alternative to phage-and yeast-display technologies to improve the breadth, potency, or bioavailability of existing antibody or even another biologic. Finally, this technology may form the basis of future vaccines and cell therapies, following a path established by chimeric antigen-receptor (CAR) T cells.

Nearly every reported effort to date to reprogram primary B cells utilizes a conserved intron downstream of the VDJ-recombined variable region.^9, 11-13^ A typical insertion cassette initiates with a poly(A) tail to terminate transcription of the native variable region, followed by an exogenous promoter, a human bNAb light-chain variable-region sequence, a P2A peptide or linker, and a heavy-chain variable-region sequence with a splice donor that promotes splicing to native constant genes. This approach is efficient because every mature B cell could theoretically be modified, because HDRT with long homology arms can be used, and because a single editing event introduces both variable chains at once. This efficiency makes this approach attractive to most investigators, but other approaches that preserve the organization of the heavy-chain locus have been attempted. For example, Voss et al.^10^ have explored an alternative in which the entire native variable-region was replaced by the heavy-chain variable of the bNAb PG9, before reverting to the more common intron-targeting approach.

A common property of all previous investigations is that an entire heavy chain or heavy-chain/light-chain pair are introduced. Thus initially B cells express a monoclonal antibody which then can diversify through SHM. As a consequence, one major contributor to antibody diversity, namely combinatorial diversity, is bypassed. Such an approach is necessary for many HIV-1 bNAbs because antigen recognition is distributed across multiple CDR loops. However most known V2-glycan/apex bNAbs have long, acidic HCDR3 regions that make an unusually large contribution to Env recognition. Moreover, neutralization with these antibodies is especially potent, typically ten-fold higher than with other classes of bNAbs. Finally, antibodies of this class, uniquely recognizing a quaternary epitope, are especially sensitive to the quality of HIV-1 antigens. We therefore undertook to reprograming human B-cells to express the HCDR3 of a potentV2-glycan/apex antibody while largely preserving the combinatorial diversity of the reservoir. Ultimately, we anticipate this diversity will enhance and personalize the adaptive humoral response to the diversity of HIV-1 isolates in reservoirs of infected humans.

However, before this concept can be tested, we had to address several challenges unique to introducing an HCDR3 into primary human B cells. These challenges arose from two sources. First, for optimal editing, a double-strand break should be introduced near the insert region, in this case at the 5’ of a commonly used J gene such as JH4. However, due to junctional diversity, this region is highly variable in a diverse repertoire. Second, HDRT with long homology arms are typically more efficient, but long arms would complement only a narrow BCR subsets or overwrite the native VH gene.

To address the first challenge, we initiated studies with the CRISPR effector protein Cas12a. We began with Cas12a because, unlike the more commonly employed Cas9, this CRISPR effector protein cleaves distally from its PAM and seed regions. Thus a more variable region can be cleaved from a more predictable gRNA target sequence. However, most commonly studied Cas12a orthologs, including LbCas12a and AsCas12a, use restrictive PAM recognition sequences^31^ absent in human JH4 sequences. We therefore characterized a number of less studied Cas12a variants, and identified one, MbCas12a, that efficiently recognized a GTTC PAM present at an optimal location in both human and rodent JH4 genes. Thus electroporated Mb2Cas12a RNP efficiently introduced double-strand breaks near the 5’ of the JH4-encoded region in Jeko-1 cells and in primary human B cells.

Our second challenge arose from the unpredictability of the VH-encoded region in the diverse repertoire of primary B cells, precluding the use of long HDRT. We accordingly designed HDRT that recognized short consensus sequences at the 3’ of three VH-gene families. We also optimized the efficiency of editing using these shorter HDRT. To do so, we first evaluated the impact of two parameters that have been proposed to alter editing efficiency. First, we asked whether the HDRT should complement the coding or non-coding sequence, and we also investigated whether it should complement the gRNA target strand or its opposite. Both parameters have be reported to contribute to editing efficiencies of other systems, but neither of these variables had a dramatic impact on editing efficiency in our study. Further analysis identified a distinct, decisive factor in editing efficiency, namely the length the 3’ mismatch tail. While 5’-3’ resection is among the first events in HDR, removal of 3’ end is rate limiting. Our data suggest that the pace of removal of the 3’ end of the HDRT complementary strand is especially critical, and if the necessary deletion is greater than 10 nucleotides, editing is significantly impaired. These data are consistent with previous observations that Pol δ, the polymerase responsible for 3’ extension of the HDRT-associated strand, has modest 3’ exonuclease activity.^47, 48^ Regardless of the underlying mechanism, this optimization enabled editing efficiencies with Mb2Cas12a and short single-stranded HDRT comparable to those reported with Cas9 and much longer HDRT.

With these tools in hand, we showed that the HCDR3 regions of primary human B cells could be reprogrammed to encoded three novel sequences, two of which derived from HIV-1 bNAbs. In each case edited cells could be enriched by FACS with an appropriate antigen while still largely retaining the diversity of the edited repertoire. Interestingly, in the case of cells edited to express the PG9 HCDR, this approach more efficiently enriched BCR encoded by the V3-30 heavy chain than for V3-33, the heavy-chain gene from which the bNAb PG9 originally derived. We confirmed this observation by showing that a PG9 variant constructed from this germline form of this VH gene neutralizes more efficiently than one constructed from the germline V3-33 gene. As importantly, a number of VH genes enriched in this manner bound SOSIP trimers and neutralized HIV-1. Thus, at least in the case of PG9, a range of modified BCR could respond to a SOSIP antigen or to HIV-1 emerging from a reactivated reservoir.

In short, we have overcome several challenges associated with introducing an exogenous HCDR3 sequence into a diverse repertoire of human BCR. In the process we have identified and characterized a Cas12a ortholog especially useful introduced double-strand breaks near the DJ junction of a recombined heavy-chain, and described an optimized approach for replacing genomic regions with short HDRT. Finally, we showed that this approach could create a diverse repertoire of B cells capable of recognizing a critical epitope of HIV-1 Env. These studies establish foundations for proof-of-concept studies in primate models of HIV-1 infection.

## MATERIAL AND METHODS

### Plasmids

Wild-type Mb2Cas12a (pcDNA3.1-hMb2Cpf1), Mb3Cas12a (pcDNA3.1-hMb3Cpf1), TsCas12a (pcDNA3.1-hTsCpf1) and BsCas12a (pcDNA3.1-hBsCpf1) plasmids were gifts from Dr. Feng Zhang (Addgene plasmid numbers 69982, 69988, 69983, and 42230, respectively). pMAL-his-LbCpf1-EC was a gift from Dr. Jin-Soo Kim (Addgene plasmid number 79008) and was used to express Mb2Cas12a in *E. coli* for protein production. For Cas12a protein production, each Cas12a gene was codon-optimized for E. coli, synthesized by IDT, and cloned into pMAL-his-LbCpf1-EC vector.

### Mb2Cas12a protein production and purification

Expression cassette of maltose binding protein (MBP)-Mb2Cas12a-His in pMal vector was transformed to Rosetta 2 (DE3) (Novagen) competent cells. Single colony was first grown in 5 mL, then scaled up to 10L for production, in LB broth with 4 μm/mL chloramphenicol and 100 ug/mL carbenicillin. Cell cultures were grown to OD ∼0.5 before placing on ice for 15 min, and added with 0.5 mM IPTG at 16°C to induce expression. After 18hr of incubation, cells were resuspended in the buffer with 50 mM NaH2PO4, 500 mM NaCl, 15 mM imidazole, 10% glycerol, and 10 mM Tris at pH 8.0, then sonicated on ice for 20 min at 18 W output before clarified by centrifugation for 25 min at 50,000 g. Clarified supernatant was loaded to the HisTrap FF column (GE Healthcare) and eluted with linear imidazole gradient from 10 mM to 300 mM using ÄKTA explorer (GE Healthcare). To remove the N-terminal MBP tag, the protein elution fractions were pooled and concentrated with 50 kDa molecular weight cutoff ultrafiltration unit (Millipore), then 1 mg of TEV protease was used per 50 mg protein forcleavage during dialyzing to the buffer with 250 mM NaCl, 0.5 mM EDTA, 1 mM DTT, and 20 mM HEPES with pH 7.4 for 48 h at 4°C. For cation exchange chromatography, the protein was diluted with 2-fold volume of 20 mM HEPES with pH 7.0 and loaded on HiTrap SP HP column (GE Healthcare) equilibrated with 100 mM NaCl, 20 mM HEPES at pH 7.0. Proteins were eluted with a linear NaCl gradient from 100 mM to 2 M, then further purified by size exclusion chromatography with Superdex 200 26/60 column (GE Healthcare) with the protein storage buffer (500 mM NaCl, 0.1 mM EDTA, 1 mM DTT, 10% glycerol and 20 mM HEPES at pH 7.5). Pure protein fractions were pooled and concentrated followed with endotoxin removal with columns from Pierce.

### RNP formation and electroporation

Mb2Cas12a, AsCas12a and SpCas9 gRNAs were ordered from IDT. RNAs were resuspended in Rnase-free water and refolded by incubation at 95°C for 5 min and cooling down at room temperature for 1 h. For each electroporation sample, RNP complexes were formed by mixing 200 pmol of Mb2Cas12a, AsCas12a (Alt-R® A.s. Cas12a (Cpf1) V3 from IDT) or SpCas9 (Alt-R S.p. Cas9 Nuclease V3 from IDT) with 300 pmol of crRNA and PBS. The RNP mixture was incubated at room temperature for 15 min to 30 min, then added with 600 pmol of ssDNA HDRT. For the mixture of HDRT (VH1, VH3, and VH4) to target primary cells, 200 pml of each was used. Jeko-1 cells (2 million/sample) and human primary B cells (4 million/sample) were harvested and rinsed with PBS before resuspension in electroporation solution. Cells were electroporated using Lonza 4D modules according to Lonza’s protocols. After electroporation, cells were incubated in the cuvette for 15 minutes at room temperature before transferring to the antibiotics-free media. Culture media was refreshed in 24 hours.

### HIV protein and antibodies

BG505 SOSIP v5.2 ds (E64K A316W A73C-A561C I201C-A433C)^41^ and BG505 E2p^43^ were constructed as previously described. Amino acid sequences were codon optimized and synthesized by IDT and cloned into the CMV/R expression plasmid following a human IGH signal peptide. The apex negative mutant (dBG505) was constructed by altering the V2 basic patch of the BG505 apex from RDKKQK to IDNVQQ to abolish PGT145 and PG9 binding. All proteins were produced in transiently transfected Expi293F (Invitrogen) cells. Protein constructs were co-transfected with plasmids encoding furin, FGE (formylglycine generating enzyme), and PDI (protein disulfide isomerase), respectively (at 4:1:1:1 ratio) using FectoPRO (Polyplus) according to the manufacturer’s protocol. Supernatants were harvested 5 days after transfection, filtered, and purified with CH01 or PGT145 affinity column. Proteins were eluted with gentle Ag/Ab elution buffer (21027, Thermo). The elution was exchanged to buffer (358 mM HEPES, 75 mM NaCl pH 8.0). For antibody production, heavy and light chain plasmids were co-transfected (1:1.25 ratio) in Expi293F cells. The supernatants were harvested five days later, and the IgG was purified using protein A Sepharose (GE Healthcare) and eluted with gentle Ag/Ab elution buffer (21027, Thermo), following buffer-exchanged into PBS.

### Human cell culture

Human blood samples were obtained through OneBlood. Peripheral blood mononuclear cells were isolated by density gradient centrifugation with Ficoll (GE Healthcare), stored in liquid nitrogen, then thawed in a 37°C water bath and resuspended in human B cell medium composed of RPMI-1640 with GlutaMAX, supplemented with 10% FBS or human serum, 10 mM HEPES, 1 mM sodium pyruvate, and 53 µM 2-mercaptoethanol (Gibco). B cells were isolated by magnetic sorting using the Human B Cell Isolation Kit II (130-091-151) from Miltenyi according to the manufacturer’s instructions and cultured in the above medium supplemented with 2 µg/ml anti-human RP105 antibody clone (312907, BioLegend).

### Flow cytometry and cell sorting

IgM expression was detected by FITC anti-human IgM antibody (clone MHM-88, Biolegend). HA insertion was detected by APC anti-HA antibody (clone 16B12, Biolegend). Reprogrammed specificity of B cells were validated by binding with PSG2, SOSIP, E2p proteins conjugated with different fluorophores using Lightning-Link Antibody Labeling Kits according to manufacturer’s instructions. Cultured cells were harvested by centrifugation and rinsed by FACS buffer (PBS, 0.5% BSA, 1 mM EDTA) and resuspend to 10 million/mL in 100 μ*l*. Cells were stained for 20 min on ice with fluorescently labeled antibodies (1 μg/mL) or proteins (3 μg/mL), then washed for twice with FACS buffer before measuring fluorescence by BD Accuri C6 flow cytometer or sorting by BD FACSAria Fusion sorter. Flow data were analyzed by FlowJo software.

### Sequencing and analysis of the edited B cell IgH repertoire

Human B cells harvested post gene-editing or cell sorting were lysed for RNA extraction by the RNeasy Micro Kit (74004, Qiagen). Primers used for reverse transcription and library amplification were modified from previous^29^. First-strand cDNA synthesis was performed on 11 μl of total RNA using 10 pmol of each primer in a 20ul total reaction (SuperScript III, Thermo Fisher) using the manufacturer’s protocol. Residual primers and dNTPs were degraded enzymatically (ExoSAP-IT, Thermo Fisher) according to the manufacturer’s protocol. Second-strand synthesis reaction was carried out in 100 μl using 10 pmol of each primer (HotStarTaq Plus, Qiagen). Residual primers and dNTPs were again degraded enzymatically (ExoSAP-IT) and dsDNA was purified using 0.8 volumes of SPRI beads (SPRIselect, Beckman Coulter Genomics). 40uL of eluted dsDNA was amplified again with 10 pmol of each primer in a 100 μl total reaction volume (HotStarTaq Plus). DNA was purified from the PCR reaction product using 0.8 volumes of SPRI beads (SPRIselect). 10 μl of the eluted PCR product was used in a final indexing using NEBNext Multiplex Oligos for Illumina (E7710S, NEB) following the manufacturer’s instruction. PCR products were purified with 0.7 volumes of SPRI beads (SPRIselect). SPRI-purified libraries were sequenced on an Illumina MiSeq using 2×250 bp. Sequencing reads were processed and analyzed. Briefly, paired reads were merged with PANDAseq^49^ using the default merging algorithm then trimmed and collapsed by UMI through Migec using the “checkout” algorithm. Processed reads were mapped by MiGMAP based on IgBlast^50, 51^.

### Neutralization assay

Pseudoviruses were produced by co-transfection of different HIV envelope plasmids acquired through the NIH AIDS Reagents Program along with NL4-3-ΔEnv in HEK293T cells using PEIpro (Polyplus). Supernatant was harvested 48h post transfection, clarified by centrifugation and 0.45 µm filter, and aliquoted for storage at -80°C. TZM-bl neutralization assays were performed as previously described^52^. Briefly, titrated antibodies in 96-well plates were incubated with pseudotyped viruses at 37°C for 1 hour. TZM-bl cells were then added to the wells with 50,000 cells/well. Cells were then incubated for 48 hours at 37°C. At 48h post infection, cells were lysed in wells and subjected to Firefly luciferase assays. Viral entry was determined using Britelite Plus (PerkinElmer), and luciferase expression was measured using a Victor X3 plate reader (PerkinElmer).

### Statistical analysis

Data expressed as mean values ± SD or SEM, and all statistical analysis was performed in GraphPad Prism 7.0 software. IC_50_ of antibody neutralization was analyzed using default settings for log(inhibitor) vs. normalized response method. Statistical difference was determined using non-paired Student’s t-test or one-way ANOVA with Tukey’s test. Differences were considered significant at P < 0.05.

## ACKNOWLEGEMENTS

This work is supported by NIH R37 AI091476 and DP1 DA043912 (PI: Farzan). The authors would like to thank Xiaohua Wu, Ph.D. for scientific advice on DNA repair mechanisms, and Hyeryun Choe, Ph.D for reviewing of the manuscript. The authors declare they have no financial interests.

## AUTHOR CONTRIBUTIONS

T.O., W.H., G.Z., and M.F. conceived of this study. G.Z. and M.F. provided guidance for experimental design. T.O., W.H., B.Q., Y.G., P.K., H.P., M.D.G., M.H.T., Y.Y., X.Z., and H.W. performed all experiments. T.O., W.H., and M.F. wrote the manuscript.

## Reference

1. West Jr AP, Scharf L, Scheid JF, Klein F, Bjorkman PJ, Nussenzweig MC. Structural insights on the role of antibodies in HIV-1 vaccine and therapy. Cell. 2014;156(4):633–48.

2. Mascola JR, Haynes BF. HIV‐1 neutralizing antibodies: understanding nature’s pathways. Immunological reviews. 2013;254(1):225–44.

3. Burton DR, Hangartner L. Broadly neutralizing antibodies to HIV and their role in vaccine design. Annual review of immunology. 2016;34:635–59.

4. Escolano A, Dosenovic P, Nussenzweig MC. Progress toward active or passive HIV-1 vaccination. Journal of Experimental Medicine. 2017;214(1):3–16.

5. Jardine JG, Kulp DW, Havenar-Daughton C, Sarkar A, Briney B, Sok D, Sesterhenn F, Ereño-Orbea J, Kalyuzhniy O, Deresa I. HIV-1 broadly neutralizing antibody precursor B cells revealed by germline-targeting immunogen. Science. 2016;351(6280):1458–63.

6. Steichen JM, Lin Y-C, Havenar-Daughton C, Pecetta S, Ozorowski G, Willis JR, Toy L, Sok D, Liguori A, Kratochvil S. A generalized HIV vaccine design strategy for priming of broadly neutralizing antibody responses. Science. 2019;366(6470).

7. Klein F, Diskin R, Scheid JF, Gaebler C, Mouquet H, Georgiev IS, Pancera M, Zhou T, Incesu R-B, Fu BZ. Somatic mutations of the immunoglobulin framework are generally required for broad and potent HIV-1 neutralization. Cell. 2013;153(1):126–38.

8. Walker LM, Huber M, Doores KJ, Falkowska E, Pejchal R, Julien J-P, Wang S-K, Ramos A, Chan-Hui P-Y, Moyle M. Broad neutralization coverage of HIV by multiple highly potent antibodies. Nature. 2011;477(7365):466–70.

9. Huang D, Tran JT, Olson A, Vollbrecht T, Tenuta M, Guryleva MV, Fuller RP, Schiffner T, Abadejos JR, Couvrette L. Vaccine elicitation of HIV broadly neutralizing antibodies from engineered B cells. Nature communications. 2020;11(1):1–10.

10. Voss JE, Gonzalez-Martin A, Andrabi R, Fuller RP, Murrell B, McCoy LE, Porter K, Huang D, Li W, Sok D. Reprogramming the antigen specificity of B cells using genome-editing technologies. Elife. 2019;8:e42995.

11. Hartweger H, McGuire AT, Horning M, Taylor JJ, Dosenovic P, Yost D, Gazumyan A, Seaman MS, Stamatatos L, Jankovic M. HIV-specific humoral immune responses by CRISPR/Cas9-edited B cells. Journal of Experimental Medicine. 2019;216(6):1301–10.

12. Moffett HF, Harms CK, Fitzpatrick KS, Tooley MR, Boonyaratanakornkit J, Taylor JJ. B cells engineered to express pathogen-specific antibodies protect against infection. Science immunology. 2019;4(35).

13. Nahmad AD, Raviv Y, Horovitz-Fried M, Sofer I, Akriv T, Nataf D, Dotan I, Carmi Y, Burstein D, Wine Y. Engineered B cells expressing an anti-HIV antibody enable memory retention, isotype switching and clonal expansion. Nature communications. 2020;11(1):1–10.

14. Giudicelli V, Duroux P, Ginestoux C, Folch G, Jabado-Michaloud J, Chaume D, Lefranc M-P. IMGT/LIGM-DB, the IMGT® comprehensive database of immunoglobulin and T cell receptor nucleotide sequences. Nucleic acids research. 2006;34(Suppl_1):D781–D4.

15. Rolink A, Melchers F. Molecular and cellular origins of B lymphocyte diversity. Cell. 1991;66(6):1081–94.

16. Elhanati Y, Sethna Z, Marcou Q, Callan Jr CG, Mora T, Walczak AM. Inferring processes underlying B-cell repertoire diversity. Philosophical Transactions of the Royal Society B: Biological Sciences. 2015;370(1676):20140243.

17. Schatz DG, Ji Y. Recombination centres and the orchestration of V (D) J recombination. Nature Reviews Immunology. 2011;11(4):251–63.

18. Bassing CH, Swat W, Alt FW. The mechanism and regulation of chromosomal V (D) J recombination. Cell. 2002;109(2):S45–S55.

19. Mesin L, Ersching J, Victora GD. Germinal center B cell dynamics. Immunity. 2016;45(3):471–82.

20. Leslie A, Pfafferott K, Chetty P, Draenert R, Addo M, Feeney M, Tang Y, Holmes E, Allen T, Prado J. HIV evolution: CTL escape mutation and reversion after transmission. Nature medicine. 2004;10(3):282–9.

21. Abbott RK, Lee JH, Menis S, Skog P, Rossi M, Ota T, Kulp DW, Bhullar D, Kalyuzhniy O, Havenar-Daughton C. Precursor frequency and affinity determine B cell competitive fitness in germinal centers, tested with germline-targeting HIV vaccine immunogens. Immunity. 2018;48(1):133-46. e6.

22. Dosenovic P, Kara EE, Pettersson A-K, McGuire AT, Gray M, Hartweger H, Thientosapol ES, Stamatatos L, Nussenzweig MC. Anti–HIV-1 B cell responses are dependent on B cell precursor frequency and antigen-binding affinity. Proceedings of the National Academy of Sciences. 2018;115(18):4743–8.

23. Andrabi R, Voss JE, Liang C-H, Briney B, McCoy LE, Wu C-Y, Wong C-H, Poignard P, Burton DR. Identification of common features in prototype broadly neutralizing antibodies to HIV envelope V2 apex to facilitate vaccine design. Immunity. 2015;43(5):959–73.

24. Lee JH, Andrabi R, Su C-Y, Yasmeen A, Julien J-P, Kong L, Wu NC, McBride R, Sok D, Pauthner M. A broadly neutralizing antibody targets the dynamic HIV envelope trimer apex via a long, rigidified, and anionic β-hairpin structure. Immunity. 2017;46(4):690–702.

25. Zetsche B, Strecker J, Abudayyeh OO, Gootenberg JS, Scott DA, Zhang F. A Survey of Genome Editing Activity for 16 Cpf1 orthologs. bioRxiv. 2017:134015.

26. Doudna JA, Charpentier E. The new frontier of genome engineering with CRISPR-Cas9. Science. 2014;346(6213).

27. Wang Y, Liu KI, Sutrisnoh N-AB, Srinivasan H, Zhang J, Li J, Zhang F, Lalith CRJ, Xing H, Shanmugam R. Systematic evaluation of CRISPR-Cas systems reveals design principles for genome editing in human cells. Genome biology. 2018;19(1):1–16.

28. Liang X, Potter J, Kumar S, Ravinder N, Chesnut JD. Enhanced CRISPR/Cas9-mediated precise genome editing by improved design and delivery of gRNA, Cas9 nuclease, and donor DNA. Journal of biotechnology. 2017;241:136–46.

29. Briney B, Inderbitzin A, Joyce C, Burton DR. Commonality despite exceptional diversity in the baseline human antibody repertoire. Nature. 2019;566(7744):393–7.

30. Volpe JM, Kepler TB. Large-scale analysis of human heavy chain V (D) J recombination patterns. Immunome research. 2008;4(1):1–10.

31. Zetsche B, Gootenberg JS, Abudayyeh OO, Slaymaker IM, Makarova KS, Essletzbichler P, Volz SE, Joung J, Van Der Oost J, Regev A. Cpf1 is a single RNA-guided endonuclease of a class 2 CRISPR-Cas system. Cell. 2015;163(3):759–71.

32. Jinek M, Chylinski K, Fonfara I, Hauer M, Doudna JA, Charpentier E. A programmable dual-RNA–guided DNA endonuclease in adaptive bacterial immunity. science. 2012;337(6096):816–21.

33. Yamano T, Zetsche B, Ishitani R, Zhang F, Nishimasu H, Nureki O. Structural basis for the canonical and non-canonical PAM recognition by CRISPR-Cpf1. Molecular cell. 2017;67(4):633-45. e3.

34. Shalem O, Sanjana NE, Hartenian E, Shi X, Scott DA, Mikkelsen TS, Heckl D, Ebert BL, Root DE, Doench JG. Genome-scale CRISPR-Cas9 knockout screening in human cells. Science. 2014;343(6166):84–7.

35. Hur JK, Kim K, Been KW, Baek G, Ye S, Hur JW, Ryu S-M, Lee YS, Kim J-S. Targeted mutagenesis in mice by electroporation of Cpf1 ribonucleoproteins. Nature biotechnology. 2016;34(8):807–8.

36. Paix A, Folkmann A, Goldman DH, Kulaga H, Grzelak MJ, Rasoloson D, Paidemarry S, Green R, Reed RR, Seydoux G. Precision genome editing using synthesis-dependent repair of Cas9-induced DNA breaks. Proceedings of the National Academy of Sciences. 2017;114(50):E10745–E54.

37. Richardson CD, Ray GJ, DeWitt MA, Curie GL, Corn JE. Enhancing homology-directed genome editing by catalytically active and inactive CRISPR-Cas9 using asymmetric donor DNA. Nature biotechnology. 2016;34(3):339–44.

38. Walker LM, Phogat SK, Chan-Hui P-Y, Wagner D, Phung P, Goss JL, Wrin T, Simek MD, Fling S, Mitcham JL. Broad and potent neutralizing antibodies from an African donor reveal a new HIV-1 vaccine target. Science. 2009;326(5950):285–9.

39. Sok D, van Gils MJ, Pauthner M, Julien J-P, Saye-Francisco KL, Hsueh J, Briney B, Lee JH, L. KM, Lee PS. Recombinant HIV envelope trimer selects for quaternary-dependent antibodies targeting the trimer apex. Proceedings of the National Academy of Sciences. 2014;111(49):17624–9.

40. Sanders RW, Derking R, Cupo A, Julien J-P, Yasmeen A, de Val N, Kim HJ, Blattner C, de la Peña AT, Korzun J. A next-generation cleaved, soluble HIV-1 Env trimer, BG505 SOSIP. 664 gp140, expresses multiple epitopes for broadly neutralizing but not non-neutralizing antibodies. PLoS pathog. 2013;9(9):e1003618.

41. de la Peña AT, Julien J-P, de Taeye SW, Garces F, Guttman M, Ozorowski G, Pritchard LK, Behrens A-J, Go EP, Burger JA. Improving the immunogenicity of native-like HIV-1 envelope trimers by hyperstabilization. Cell reports. 2017;20(8):1805–17.

42. Hoffhines AJ, Damoc E, Bridges KG, Leary JA, Moore KL. Detection and purification of tyrosine-sulfated proteins using a novel anti-sulfotyrosine monoclonal antibody. Journal of Biological Chemistry. 2006;281(49):37877–87.

43. He L, De Val N, Morris CD, Vora N, Thinnes TC, Kong L, Azadnia P, Sok D, Zhou B, Burton DR. Presenting native-like trimeric HIV-1 antigens with self-assembling nanoparticles. Nature communications. 2016;7(1):1–15.

44. Briney B, Sok D, Jardine JG, Kulp DW, Skog P, Menis S, Jacak R, Kalyuzhniy O, De Val N, Sesterhenn F. Tailored immunogens direct affinity maturation toward HIV neutralizing antibodies. Cell. 2016;166(6):1459-70. e11.

45. Escolano A, Steichen JM, Dosenovic P, Kulp DW, Golijanin J, Sok D, Freund NT, Gitlin AD, Oliveira T, Araki T. Sequential immunization elicits broadly neutralizing anti-HIV-1 antibodies in Ig knockin mice. Cell. 2016;166(6):1445-58. e12.

46. Tian M, Cheng C, Chen X, Duan H, Cheng H-L, Dao M, Sheng Z, Kimble M, Wang L, Lin S. Induction of HIV neutralizing antibody lineages in mice with diverse precursor repertoires. Cell. 2016;166(6):1471-84. e18.

47. Pâques F, Haber JE. Two pathways for removal of nonhomologous DNA ends during double-strand break repair in Saccharomyces cerevisiae. Molecular and Cellular Biology. 1997;17(11):6765–71.

48. Anand R, Beach A, Li K, Haber J. Rad51-mediated double-strand break repair and mismatch correction of divergent substrates. Nature. 2017;544(7650):377–80.

49. Masella AP, Bartram AK, Truszkowski JM, Brown DG, Neufeld JD. PANDAseq: paired-end assembler for illumina sequences. BMC bioinformatics. 2012;13(1):1–7.

50. Shugay M, Britanova OV, Merzlyak EM, Turchaninova MA, Mamedov IZ, Tuganbaev TR, Bolotin DA, Staroverov DB, Putintseva EV, Plevova K. Towards error-free profiling of immune repertoires. Nature methods. 2014;11(6):653–5.

51. Shugay M, Bagaev DV, Turchaninova MA, Bolotin DA, Britanova OV, Putintseva EV, Pogorelyy MV, Nazarov VI, Zvyagin IV, Kirgizova VI. VDJtools: unifying post-analysis of T cell receptor repertoires. PLoS computational biology. 2015;11(11):e1004503.

52. Montefiori DC. Measuring HIV neutralization in a luciferase reporter gene assay. HIV protocols: Springer; 2009. p. 395–405.

