## Supplemental Figures 1 and 2 for "Efficient reprogramming of the heavy-chain CDR3 regions of a human antibody repertoire"

**Contents:**

**Figure S1. Mb2Cas12a and SpCas9 target regions.**

Relevant to Figures 2 and 3.

**Figure S2. Consensus sequences of the 3' region human VH1, VH3, and VH4 regions.**

Relevant to Figures 5 and 6.

### SUPPLEMENTARY FIGURE LEGENDS

**Figure S1. Mb2Cas12a and SpCas9 target regions.** A graphical representation of the Jeko-1-cell heavy-chain locus with VH, DH, JH, CH, and HCDR3 regions represented in blue, yellow, purple, gray, and green, respectively. The Jeko-1-cell heavy-chain derives from VH2-70 and JH4 genes, as indicated. Black bar and arrow indicate HCDR3 region whose nucleotide and amino-acids are shown below. Four distinct Mb2Cas12a (orange) and SpCas9 (cyan) PAM and matching cleavage sites, used in Figures 2 and 3, are indicated. Note that Mb2Cas12a leaves 5' overhangs and that SpCas9 creates blunt ends. For each cut site, each DNA strand is labelled according to whether it is the target (T) or non-target (NT) strand of the indicated CRISPR protein and gRNA.

**Figure S2. Consensus sequences of the 3' region human VH1, VH3, and VH4 regions.** 5' homology arms of HDRT used to edit primary human B cells, as shown in Figures 5 and 6, were designed based on the intrafamily conservation of the 3' of the indicated VH genes, as shown. Sequence logo presents conserved residues, full height indicates conserved within the indicated family and smaller letters indicated less conservation. The 5' homology arms used in the figures are shown in black.

**Figure S1**

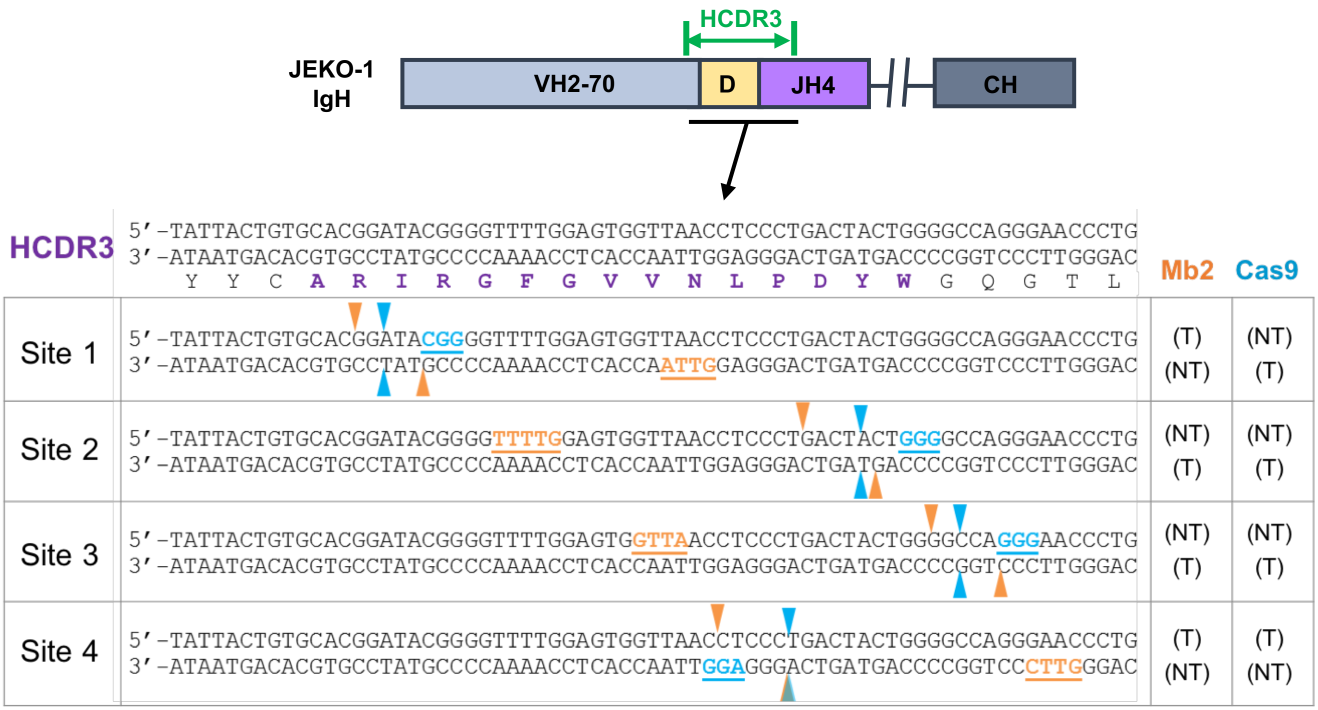
HCDR3
JEKO-1
IgH
VH2-70
D
JH4
CH
HCDR3
5'-TATTACTGTGCACGGATACGGGGTTTTGGAGTGGTTAACCTCCCTGACTACTGGGGCCAGGGAACCCTG
3'-ATAATGACACGTGCCTATGCCCCAAAACCTCACCAATTGGAGGGACTGATGACCCCGGTCCCTTGGGAC
Y Y C A R I R G F G V V N L P D Y W G Q G T L
Mb2
Cas9
Site 1
(T) (NT)
(NT) (T)
Site 2
(NT) (T)
(NT) (T)
Site 3
(NT) (T)
(NT) (T)
Site 4
(T) (NT)
(T) (NT)

### Figure S2

**Consensus sequence of VH1-FR3:**

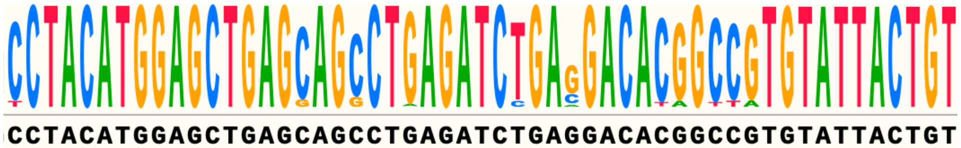


**Consensus sequence of VH3-FR3:**

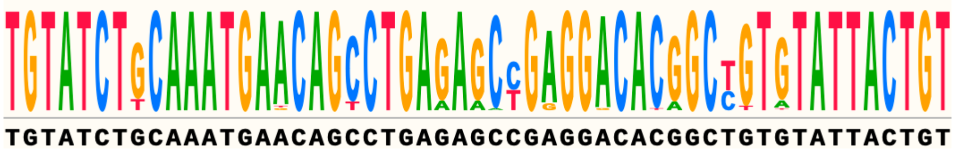


**Consensus sequence of VH4-FR3:**

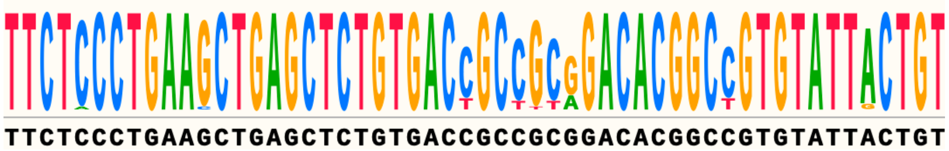
